# Diffusion MR imaging in the cortical spinal tract of idiopathic scoliosis

**DOI:** 10.1101/2022.05.31.494193

**Authors:** Edit Frankó, Olivier Joly, Olivier A. Coubard, Jean-Claude Baudrillard, Christian Morin, Dominique Rousié

## Abstract

Many studies have shown that idiopathic scoliosis is not only a deformity of the spine. It is often associated with neurological impairment without any macroscopic abnormality in the brain. In our previous diffusion MRI study, we demonstrated that children with right-thoracic idiopathic scoliosis had abnormal white matter microstructure of the crossing premotor fibres in the corpus callosum. Based on this, we hypothesized that similar microstructural changes could affect the main descending white matter tracts, the corticospinal tract.

We compared the fractional anisotropy values along the corticospinal tracts in ten patients with right-thoracic and ten patients with left-thoracic idiopathic scoliosis to 49 healthy controls.

We found abnormal left-right asymmetry of the fractional anisotropy values in scoliosis patients at the level of the pons. Whereas at upper levels the values were similar across all groups.

Our results suggest that abnormal sensorimotor integration at the level of the pons is associated with the development of idiopathic scoliosis.

## Introduction

Idiopathic scoliosis (IS) affects 2-3% of otherwise healthy children [1–3]. Biomechanical, genetic [4], hormonal, neural and metabolic changes have been reported in IS, however the cause is still unknown.

Neurological dysfunctions, such as impaired sensorimotor control, vestibular [5][6], postural [6][7] and locomotor [7,8] deficits, have often been observed in IS patients. This connection between IS and neural abnormalities is also supported by syndromes where patients with macroscopic brain malformation develop scoliosis, such as Horizontal Gaze Palsy with Progressive Scoliosis (HGPPS) [7–9]. HGPPS is caused by mutation in the ROBO3 gene, and is associated with the absence of the major crossing fibers in the pons and midbrain. These findings support the hypothesis that abnormal axonal development might play a role in the etiopathology of idiopathic scoliosis.

In our previous study, we examined the main crossing fiber tracts in the brain using diffusion MRI. We have shown abnormal fractional anisotropy (FA) values in the corpus callosum of patients with IS. Our findings suggested that even in children with no macroscopic brain abnormality, there are altered fiber tracts, fewer fibres or abnormal myelination that might be responsible for the sensorimotor, vestibular and postural impairments. Specifically, the abnormal FA values were found in the anterior body of the corpus callosum where the fibers coming from the premotor cortical areas cross the midline. This suggested that the corticospinal tracts that collect the descending fibers from the motor and premotor cortices might also be affected by this white matter microstructure abnormality.

In the current study, we examined children with right thoracic and left thoracic scoliosis and compared them to healthy controls. We hypothesized that scoliotic subjects may have microscopic abnormalities affecting the corticospinal tract. We focused on areas that were found abnormal in HGPPS and looked at the FA, its left-right asymmetry along the main white matter tracts and its association with the direction of the scoliosis.

## Methods

A cohort of twenty patients were recruited. Ten patients (age: 10-24years, mean=14years, all females and right handed) with right thoracic or right thoracic-left lumbar scoliosis from the CHRU Hospital of Lille, France. The other ten patients had left thoracic scoliosis and were recruited from Berck and Arras. Patients with congenital, juvenile or secondary scoliosis were excluded from the study.

Data from the right scoliosis patients and the controls were used in a previous study [10]. For right scoliosis patients, magnetic resonance images were acquired at 3 Tesla (Achieva, Philips Medical Systems) using parallel imaging SENSE-Head-8 channels coil at the University Hospital of Lille, without sedation or general anaesthesia. The sequences included a T1-weighted 3D acquisition, with echo time (TE) = 3.301 ms, flip angle = 98, repetition time (TR) = 7.199 ms, slice thickness = 1 mm, matrix = 256 × 256 and a DTI spin-echo echo planar image (SE-EPI) 78 axial slices, slice thickness = 2 mm, TR = 13,000 ms, flip angle = 90 degree, TE= 55 ms, matrix = 128 × 128 and along 32 isotropically distributed directions with b values of 1,000 s/mm2. The DTI acquisition was performed with isotropic resolution of 2 mm. The MR acquisitions in patients lasted for about 30 min (10 and 20 min for the T1 and the diffusion MR sequence, respectively).

### Control population

Data from 49 healthy controls (age: 10-18 years, mean=14 years, all females and right handed) were obtained from the NIH Pediatric MRI Data Repository created by the NIH MRI Study of Normal Brain Development. Data from controls were downloaded from the NDAR Pediatric Dataset website. The data included a T1-weighted MRI and tensor derived images including the FA maps which were acquired at 1.5 Tesla with 6 directions. The diffusion derived images had isotropic voxels of 2 mm. To validate the compatibility of the two datasets, we also acquired in one patient a DTI spin-echo echo planar image at 1.5T (GE Medical Systems Signa) with isotropic resolution of 2.5 mm.

### Data analysis

MR images from patients were first converted from DICOM to NIFTI format using the dcm2nii program that is distributed with MRIcron software. Then, diffusion images were processed to remove image distortion that arises from the effect of eddy currents on the EPI readout using eddy_correct function of the FDT software (FMRIB’s Diffusion Toolbox available in FSL software http://fsl.fmrib.ox.ac.uk). The FDT was also used to generate the diffusion tensor for each voxel, from which FA was derived. FA image was then thresholded (FA >0.2) to avoid voxels that are not part of the white matter tract and minimize inclusion of voxels with a high degree of partial volume effect [11].

**Figure 1-.**
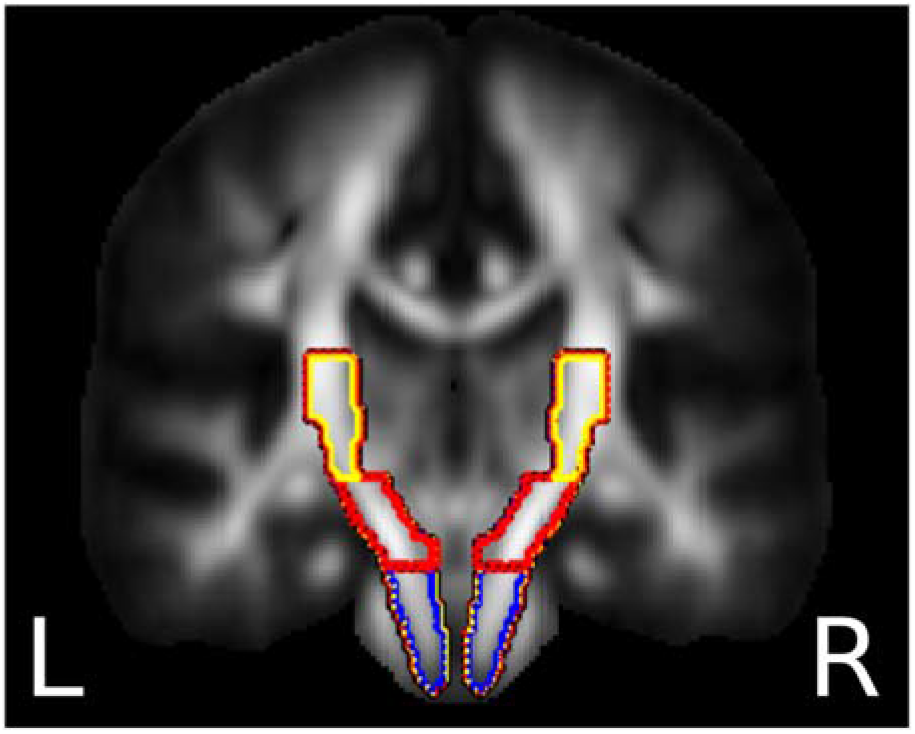
Illustration of Pons (blue), midbrain (red) and internal capsule (yellow) parts of the white matter tracts used for sampling of the individual FA maps (underlying grey scale image). Definition as provided by the JHU DTI-based white-matter atlases in FSL package [12], [13].

## Results

In the pons, the FA showed stronger left-right asymmetry in the scoliosis groups as compared to the controls, Left vs Control in Pons (t-statistic=2.634, p-value=0.0100) Right vs Control in Pons (t-statistic=3.630, p-value=0.0003). Both of the patient groups had higher FA values in the left hemisphere as compared to the right hemisphere. In the midbrain and internal capsule, we have found no significant difference in the left-right asymmetry between groups.

**Figure 2.**
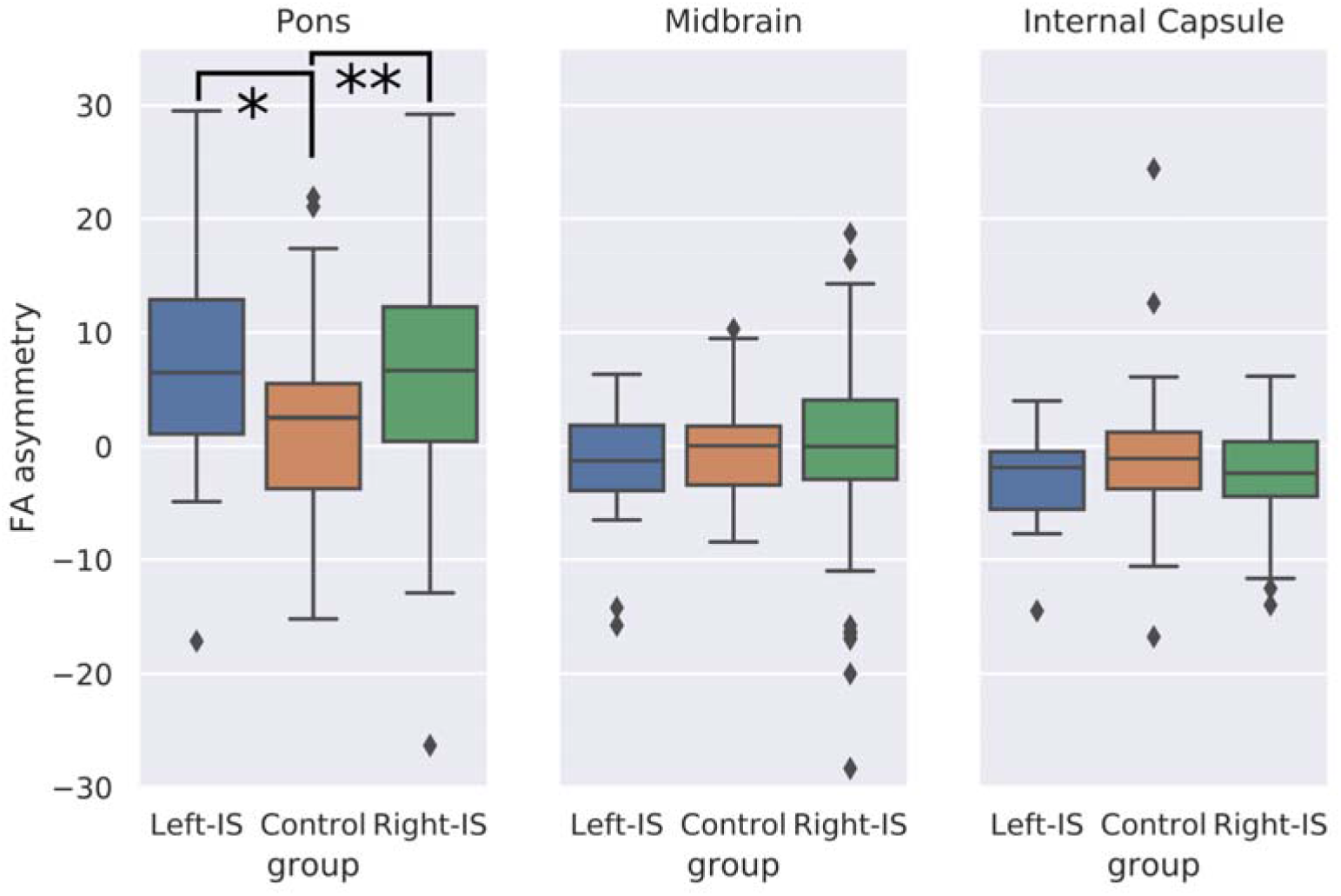
Box plot of FA asymmetry (Left hemisphere side - Right hemisphere side FA) in the 3 groups (left IS, Control and Right IS groups, in blue, red and greed) for the 3 regions Pons, Midbrain and IC. (Left FA - right FA) asymmetry.

## Discussion

In the current study, we compared the left-right asymmetry of the fractional anisotropy values of the corticospinal tract at different levels in patients with right-thoracic and left-thoracic scoliosis to healthy controls. The FA values were largely similar among the groups, and showed minimal asymmetry in the midbrain and internal capsule. However, in the pons, both patient groups showed stronger left-right asymmetry, having higher FA values in the left corticospinal tract as compared to the right tract.

Fractional anisotropy is a measurement of homogeneity of water molecule diffusion within a voxel. It is influenced by different tissue properties, such as the number and diameter of the axons, thickness of the myelin sheath and membrane permeability. Unfortunately, we cannot distinguish between these white matter microstructure changes [14,15].

Previous study by Domenech et al. [16] investigated the functional asymmetry of the motor cortex in patients with IS. Using transcranial magnetic stimulation, they found larger amplitude motor evoked potentials in right-thoracic scoliosis patients when stimulating the left hemisphere as compared to right hemisphere stimulation. This reflects relative decrease in intracortical inhibition in the motor cortex. We have found higher FA values in the left side of the pons in scoliosis patients, which can reflect increased number of fibres or thicker myelination due to increased cortical activation demonstrated by larger amplitude motor evoked potentials. This FA asymmetry in the pons could also reflect the change in fibre content of the pyramidal tract in the brainstem where the corticobulbar fibres terminate and reveal abnormalities of the corticospinal fibres.

Besides the motor cortex, the brainstem also plays an important role in motor control. As it was described in HGPPS [9], abnormality of brainstem fibre tracts is associated with the development of scoliosis. The mesopontine tegmentum and the pontomedullary reticulospinal system both play important roles in postural muscle tone and locomotion [17]. Abnormal input from the higher motor centres [18] and from the cerebellum [19,20] could cause dysfunction of the mesencaphalic-reticulospinal system, resulting in abnormal tone and locomotion, both of which have been observed in IS patients [6–8].

However, not only the motor control that influences the postural tone and the gait. Summation of sensory information from the visual, proprioceptive and vestibular receptors within the brainstem and modification of the corticospinal signal via the reticulospinal and vestibulospinal tracts are also responsible for normal posture [21]. It is difficult to examine these tracts by DTI as they are usually very small. Nonetheless, alteration of the cognitive integration of the vestibular signal with visual or somatosensory input was observed in patients with IS [22–24]. This was also supported by longer somatosensory cortical potentials in patients [25,26]. Asymmetric vestibular information was also suggested to be the underlying cause of the abnormal sensory integration. Monaural and binaural vestibular stimulation showed altered balance control in patients with AIS [27] and larger functional vestibular asymmetry [28]. This asymmetry can be explained by unilateral abnormalities of the vestibular organs as was demonstrated by Carry [29] and Rousie [30].

In this unique study where patients with left-thoracic scoliosis were compared to patients with right-thoracic scoliosis, we have found asymmetry of the descending motor pathways in patients with idiopathic scoliosis. Healthy control subjects did not show this left-right asymmetry. These results support the hypothesis that microstructural abnormalities of the motor pathways and the modification of motor control by asymmetrical sensory input can participate in the development of scoliosis.

## Acknowledgement

We are grateful to Dr Annik Delvalle and her team for their welcoming in the MRI department of Hopale Hospital in Berck and to Camille Piat and Aurore Ponchel for the imaging acquisition. We also are grateful to the team of Orthopedic surgery department of Hopale hospital for the recruitment of patients. We thank Célia Agache, MRI department of Arras hospital for her involvement in MRI registrations.

